# A putative long noncoding RNA-encoded micropeptide maintains cellular homeostasis in pancreatic β-cells

**DOI:** 10.1101/2020.05.12.091728

**Authors:** Mark Li, Fan Shao, Qingwen Qian, Wenjie Yu, Zeyuan Zhang, Biyi Chen, Dan Su, Yuwei Guo, An-Vi Phan, Long-sheng Song, Samuel B. Stephens, Julien Sebag, Yumi Imai, Ling Yang, Huojun Cao

## Abstract

Micropeptides (microproteins) encoded by transcripts previously annotated as long noncoding RNA (IncRNAs) are emerging as important mediators of fundamental biological processes in health and disease. Here we applied two computational tools to identify putative micropeptides encoded by lncRNAs that are expressed in the human pancreas. We experimentally verified one such micropeptide encoded by a β-cell- and neural cell-enriched lncRNA *TUNAR* (*also known as TUNA, HI-LNC78 or LINC00617*). We named this highly conserved 48-amino-acid micropeptide <u>B</u>eta cell- and <u>N</u>eural cell-regu<u>lin</u> (BNLN). BNLN contains a single-pass transmembrane domain and localized at the endoplasmic reticulum in pancreatic β-cells. Overexpression of BNLN lowered ER calcium levels, increased cytosolic calcium levels, and maintained ER homeostasis in response to high glucose challenge. To determine the physiological and pathological roles of BNLN, we assessed the BNLN expression in islets from mice fed with a high-fat diet and a regular diet, and found that *BNLN* is suppressed by diet-induced obesity (DIO). Conversely, overexpression of BNLN elevated glucose-stimulated insulin secretion in INS-1 cells. Lastly, BNLN overexpression enhanced insulin secretion in islets from lean and obese mice as well as from humans. Taken together, our study provides the first evidence that lncRNA-encoded micropeptides play a critical role in pancreatic β-cell function and provides a foundation for future comprehensive analyses of micropeptide function and pathophysiological impact on diabetes.

## INTRODUCTION

Large-scale projects such as ENCODE and FANTOM have found thousands of long noncoding RNAs (lncRNAs) in the human genome^1–4^. lncRNAs play an essential role in regulating islet function in health and diseases including diabetes^5^. It has been demonstrated that lncRNAs are key components of the islet regulatome, with more than 1000 lncRNAs identified in human islets and mouse β-cells^6,7^. These lncRNAs play critical roles in islet and β-cell survival, cell development, and physiological function through regulation of the islet transcriptome^5–7^. Of note, a genome-wide association study that examined T2D susceptibility loci demonstrated that dysregulation of IncRNAs is involved in pancreatic pathologies in humans^6^. In addition, dysregulation of lncRNAs has been implicated in diabetes complications, such as diabetic nephropathy and retinopathy^5,8^. Recently, several circulating lncRNAs have been identified as biomarkers of diabetes and diabetic complications^9,10^. However, although much knowledge has accumulated correlating dysregulation of lncRNA with islet pathology, the functional roles and molecular mechanisms of these lncRNAs remain poorly understood.

By definition, lncRNAs do not contain protein-coding open reading frames (ORF), which comprise a start codon, in-frame codons, and a stop codon. To accurately identify *bona fide* protein-coding ORFs, most ORF-finding computational algorithms have historically used a cutoff of 100 amino acids as the minimum size for detection. This filtering strategy dramatically reduces the likelihood of false-positive detection, but may also result in some genes that contain small open reading frames (sORFs) being mis-annotated as lncRNAs^11^. Indeed, recent studies have revealed that some of IncRNAs do contain sORFs that encode micropeptides (or microprotein, protein less than 100 amino acids in length)^12–20^. There is emerging evidence that micropeptides are important regulators in fundamental biological processes such as embryonic development^21^ and metabolism^22,23^. However, whether and how micropeptides modulate islet biology is largely unknown.

In this study, we employed two computational algorithms, PhyloCSF and RNAcode, to comprehensively survey the protein-coding potential of lncRNAs expressed in the human pancreatic islet^24,25^. Among the novel micropeptides encoding lncRNAs transcripts we identified, we characterized one putative micropeptide encoded by *TUNAR* (*also known as TUNA, HI-LNC78 or LINC00617*), which is expressed highly in human β-cells and neural cells. We then named this highly conserved micropeptide beta cell and neural cell regulin (*BNLN*). Pancreatic β-cells synthesize and secrete insulin to govern systemic glucose homeostasis^26^. We found BNLN localizes at the endoplasmic reticulum (ER) in pancreatic β-cells. Robust and highly functional endoplasmic reticulum (ER) dynamics are required for β-cells to synthesize insulin to maintain systemic glucose metabolic homeostasis^27^. The ER also plays an important role in coordinating intracellular Ca^2+^ signaling which regulates insulin secretion^28,29^. In both rodent models and humans, obesity and diabetes are characterized by ER dysfunction which leads to β-cell death and impaired islet function^30,31^. We further showed that *BNLN* is downregulated in islets from obese mice, and gain of function of *BNLN* increases glucose-induced insulin secretion in islets from obese mice as well as from human donors. These findings provide first evidence of the biological function of pancreatic micropeptides in health and disease.

## RESULTS

### An IncRNA, *TUNAR*, encodes a conserved 48-amino-acid micropeptide in the human pancreas

Recent studies have discovered that a few lncRNAs harbor small open reading frames (sORFs) that encode sORF-encoded peptides (SEPs) or micropeptides^12–20^. We hypothesized that there are additional transcripts currently annotated as lncRNAs that may encode micropeptides. To prioritize the search for potential micropeptide genes, we used two computational tools, PhyloCSF and RNAcode, to comprehensively evaluate the coding potential of lncRNAs that are expressed in human pancreatic islets^24,25^. We found several lncRNAs that are highly expressed in human pancreatic islets and contained putative sORFs (Fig. 1A). TCL1 Upstream Neural Differentiation-Associated RNA, *TUNAR (also known as TUNA, HI-LNC78 or LINC00617)* ranked at the top in both the expression level in human islets and the coding potential (Fig. 1A). Of note, its PhyloCSF and RNAcode scores were higher than many recently identified micropeptides (Fig. 1B). In addition to the pancreas, *TUNAR* is highly expressed in the brain, pituitary, testis, fallopian tube and the uterus (Supplemental Fig. 1). Knockdown of *TUNAR* has been associated with reduced insulin content and impaired glucose-stimulated insulin secretion in a human β-cell line, as well as impaired neural differentiation of mouse embryonic stem cells (mESCs)^32,33^. Genomic annotation indicated that *TUNAR* spans ~ 49.2kb and has 3 transcript isoforms: NR_132399.1, NR_132398.1 and NR_038861.1 (Fig. 1C). Further analysis of RNA-Seq read splice junctions visualized with sashimi plot revealed that NR_038861.1 is the major isoform in the pancreas (data not shown). Our computational pipeline predicted an sORF at the beginning of the last exon (which is shared among all three transcript isoforms) that is highly conserved among 100 vertebrate genomes assessed by UCSC genome browser PhyloP track (Fig. 1C). Moreover, ribosomal profiling (Ribo-seq) datasets suggest that this region is actively translated (Supplemental Fig. 2A). We then extracted other species’ DNA sequences (multiz100way) corresponding to predicted sORF and translated them to protein sequences. The predicted micropeptide sequence was highly conserved (Fig. 1D, Supplemental Fig. 3 and Supplemental Table1) and could be found in 88 species out of the 100 vertebrate species (conserved down to fishes).

**Figure 1.**
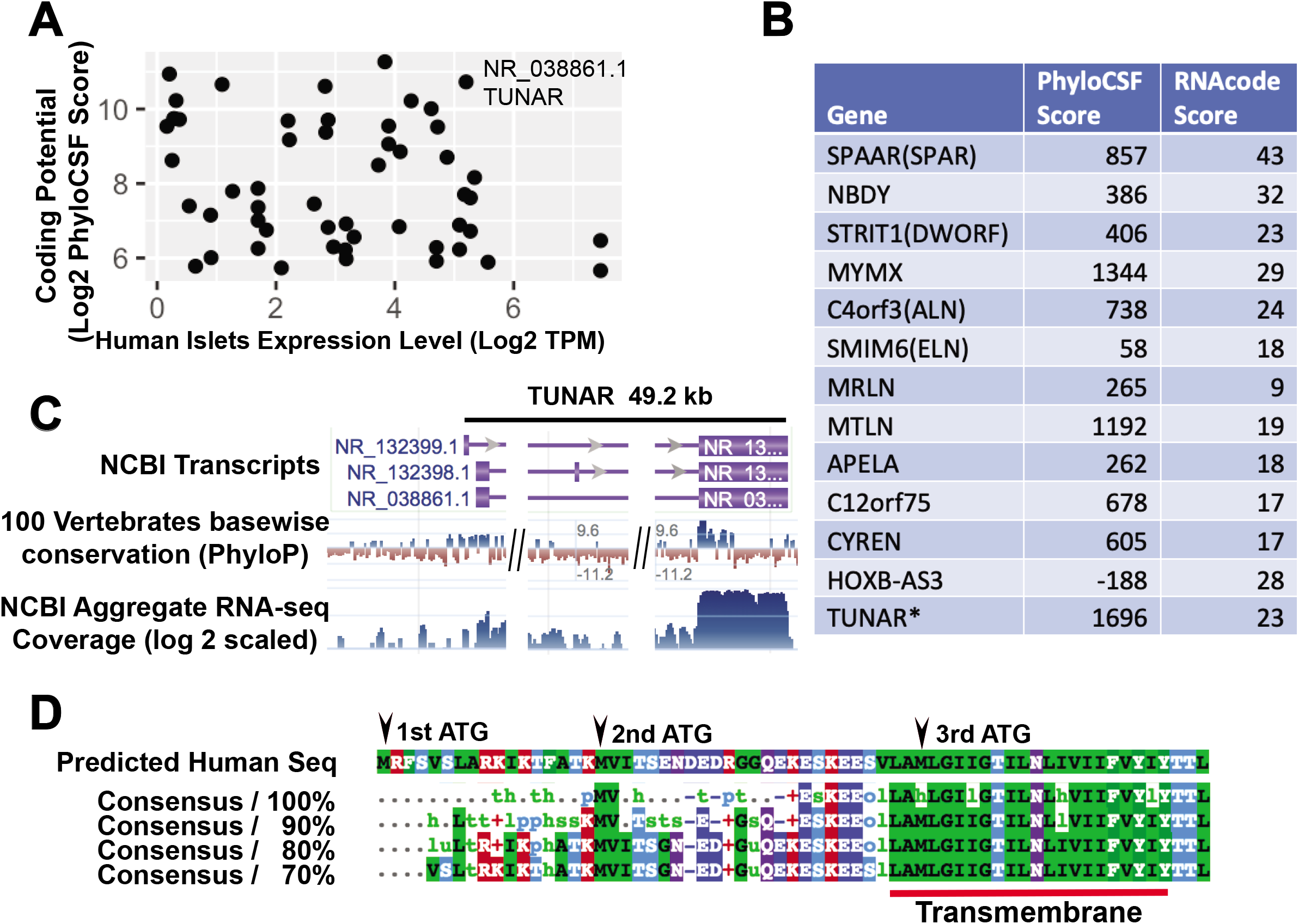
Identification of lncRNAs that have coding potential in the human pancreas. **A.** Scott plot of lncRNA expression level in human islets and coding potential. Expression level was measured by log2(TPM+1). Coding potential was measured by log2(PhyloCSF score + 1). **B.** PhyloCSF score and RNAcode score for 12 lncRNAs that have recently been found to encode micropeptides. *TUNAR* is the micropeptide encoding lncRNA identified in this study. **C.** Genomic architecture of the *TUNAR* locus. 100 vertebrates sequence conservation shows a conserved region at the beginning of last exon. **D.** Multi-species alignment of predicted *TUNAR* encoding micropeptide BNLN. A conserved transmembrane domain is located at the C-terminus. Three potential start codons were presented.

To experimentally validate the translation of the predicted sORF within *TUNAR*, we inserted three reporters (GFP, hRluc and FLAG) CDS without ATG start codon either in-frame or out-of-frame before the stop codon of the full length *TUNAR* transcript (NR_038861.1; Fig. 2A). As expected, GFP and Renilla luciferase were translated when inserted in-frame (Fig.2B&C). However, no translation was observed when GFP or hRluc were inserted out-of-frame by deletion of one or two bases between the predicted sORF and GFP/hRluc (Fig. 2B&C). Consistent with these results, western blot analysis showed a band ~ 10-kDa only when FLAG was inserted in-frame (Fig. 2D). There are 3 potential ATG initiating codons in-frame with the predicted reading frame in *TUNAR* (Fig. 1D). Multispecies alignment showed that more than half of all the species contain the 1^st^ ATG, all 88 species have the 2^nd^, and 87 species have 3^rd^ ATG of *TUNAR* (Supplemental Fig. 3). Therefore, we generated ATG deletion constructs for each of these potential initiating codons and found that removing the 2^nd^ ATG abolished translation of *TUNAR* (Fig. 2E&F). Hence, *TUNAR* contains an sORF from the 2^nd^ ATG start codon to the stop codon that translates to a 48-amino-acid micropeptide. To keep consistency with the previous nomenclature, we named this micropeptide <u>B</u>eta cell- and <u>N</u>eural cell-regu<u>lin</u>_(BNLN).

**Figure 2.**
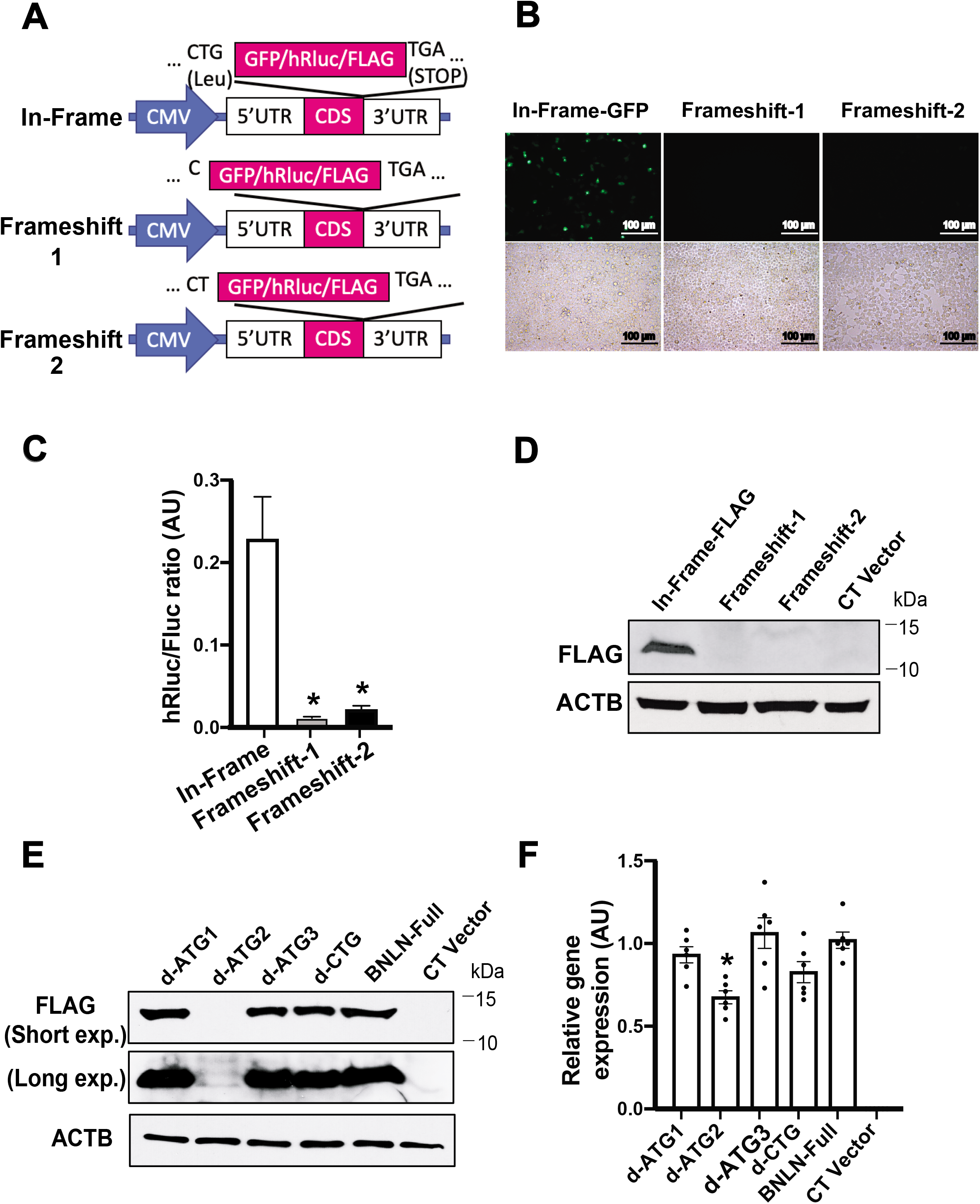
*TUNAR* encodes a novel micropeptide, BNLN. **A**. Schematic representation of *BNLN* reporter constructs. *BNLN* coding sequence (CDS) is shown in pink block. **B.** Representative images (20X) and **C.** Activity of Renilla luciferase in HEK 293T cells transfected with the indicated BNLN constructs as in (A) for 48 hr. Scale bar: 50μm. The data were normalized to firefly luciferase. * indicates statistical significance compared to in-frame constructs (n=4, biological replicates). AU, arbitrary units. **D.** Representative western blots of BNLN expression in HEK 293T cells transfected with the indicated *BNLN* constructs. CT: non-transfected cell control. **E.** Representative western blots of BNLN expression in cells transfected with the indicated *BNLN* constructs for 48hrs. **F.** Levels of mRNAs encoding for BNLN in HEK293T cells expressing full length, ATG-deleted and CTG-deleted *BNLN* CDS. The data were normalized to *ACTB* expression level. * indicates statistical significance compared to the full *BNLN* expressing cells (n=6, biological replicates). Data are shown as means ± SEM, statistical significance was determined by ANOVA followed by Sidak’s multiple comparisons test in C&F. p<0.05.

### BNLN localizes at the endoplasmic reticulum (ER) in pancreatic β cells

*BNLN/ TUNAR* is highly expressed in the human islet (Fig. 2A). *In situ* hybridization analysis in islets from human donors showed that *BNLN* expression is enriched in the insulin producing β-cells (Fig. 3A), which is in agreement with the analysis of previous RNA-Seq data^32,34^. Our *in silico* sequence analysis revealed that BNLN harbors a single-pass transmembrane domain at its C-terminus (Fig. 1D, Supplemental Fig. 2B). The multispecies alignment showed that this transmembrane domain is extremely conserved and presented in all 88 species (Fig. 1D, Supplemental Fig. 3). To determine the sub-cellular localization of BNLN, we generated N-terminal and C-terminal GFP-FLAG tagged *BNLN* constructs (Fig. 3B), then performed immunofluorescence analysis in INS-1 cells. As shown in figure 3C, in contrast to the GFP control, which was dispersed throughout cells, BNLN mainly localized at the endoplasmic reticulum (ER) in the INS-1 cell. Much weaker signal was detected in lysosomes, mitochondria, peroxisomes, or Golgi in INS-1 cells (Fig. 3D). To further assess whether BNLN is secreted or expressed extracellularly, we generated a BNLN construct fused with HiBiT (Fig. 3E). In the Nano-Glo HiBiT detection system, HiBiT-tagged protein is recognized by the complementing polypeptide LgBiT, the bound complex then reconstitutes a luminescent signal. As shown in figure 3F, BNLN mainly localized intracellularly in INS-1 cells. Taken together, these data show that BNLN is primarily located at the ER of pancreatic β-cells.

**Figure 3.**
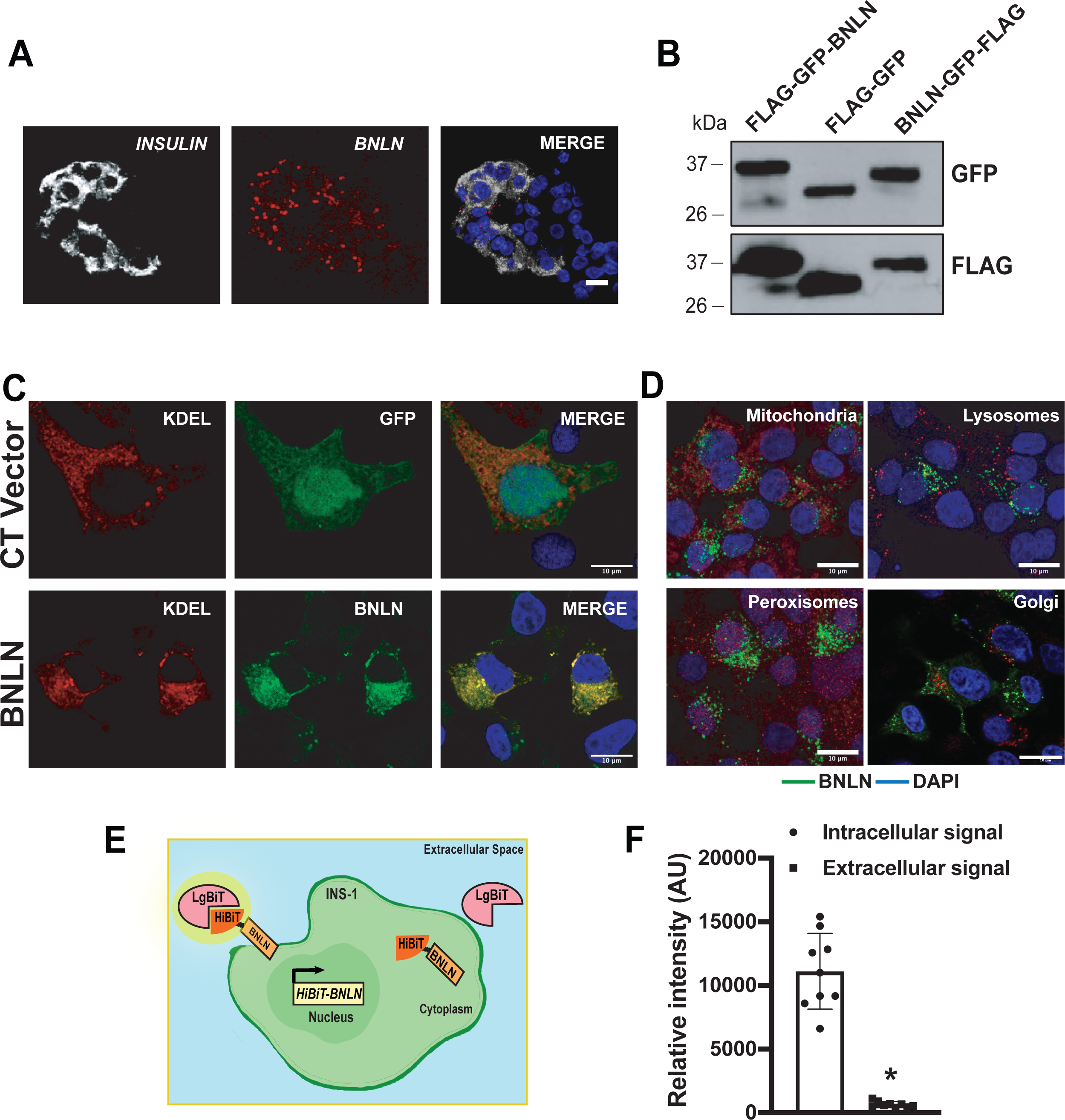
BNLN localizes at the endoplasmic reticulum in β-cells. **A.** Representative confocal images (60X) of *TUNAR* in human islets. Small molecular *in situ* hybridization of *BNLN*, and *INSULIN* (β-cell marker). Scale bar: 10μm. **B.** Representative western blots of BNLN expression in INS-1 cells transfected with the indicated *BNLN* constructs for 48hrs. **C.** Representative confocal images (63X) of BNLN in INS-1 cells co-transfected with FLAG-GFP-BNLN and KDEL-RFP (24hrs post transfection). CT vevtor: FLAG-GFP construct. Scale bar: 10μm. **D**. Representative confocal images (63X) of BNLN in INS-1 cells transfected with BNLN-GFP and stained with lysotracker (lysosomes), mitotracker (mitochondria), Acaa1 antibody (peroxisomes), and TGN38 antibody (Golgi). Scale bar: 10μm. **E**. Model of HiBiT protein tagging system in monitoring BNLN localization. INS-1 cells were transfected with HiBiT-tagged BNLN, the cell surface HiBiT was recognized by recombinant LgBiT and NanoBiT substrate in live cells. **F**. Expression of cell surface HiBiT-BNLN level in INS-1 cells 24hrs after transfecting with the HiBiT-BNLN construct. The level of cell surface HiBiT-BNLN was measured by using LgBiT and the extracellular substrate, and normalized to the total expression of the construct by lysing the cells with the lytic substrate in the presence of LgBiT. Data are shown as means ± SEM, and statistical significance was determined by Student’s t-test (p<0.05, n=3, biological replicates). AU, arbitrary units.

### BNLN modulates ER calcium homeostasis and ER function in the β cell

ER calcium homeostasis has a pivotal role in insulin production and secretion in β-cells^26^. To determine the functional effect of BNLN in the ER, we first assessed the ER Ca^2+^ ([Ca^2+^]^ER^) level by using an ER Ca^2+^ biosensor (ER-GCaMP6-150) in INS-1 cells overexpressing BNLN or the control construct. Overexpression of BNLN lowered [Ca^2+^]^ER^ level at both the basal (2.5mM) and high glucose (17mM) conditions (Fig. 4A&B), indicating a potential role of BNLN in modulating ER calcium influx. Glucose-stimulated β-cells display oscillations of the interrelated cytosolic Ca^2+^ ([Ca^2+^]^c^) and [Ca^2+^]^ER^. To evaluate the influence of the BNLN on [Ca^2+^]^c^ homeostasis, we generated FLAG- or V5-tagged *BNLN* constructs and measured [Ca^2+^]^c^ by a FLIPR Tetra system. Overexpression of BNLN increased [Ca^2+^]^c^ in INS-1 cells under both basal and high glucose conditions (Fig. 4C&D). We observed this effect with both FLAG- and V5-tagged BNLN. To further assess the effect of BNLN on depolarization-induced calcium dynamics, we measured [Ca^2+^]^c^ in INS-1 in the absence or presence of KCl. As shown in figure 4E&F while BNLN overexpression increased high glucose-induced [Ca^2+^]^c^, it did not alter KCl-mediated [Ca^2+^]^c^ flux, indicating a regulatory role of BNLN independent from depolarization dependent [Ca^2+^]^c^ flux that is primarily driven by voltage-dependent calcium channel.

**Figure 4.**
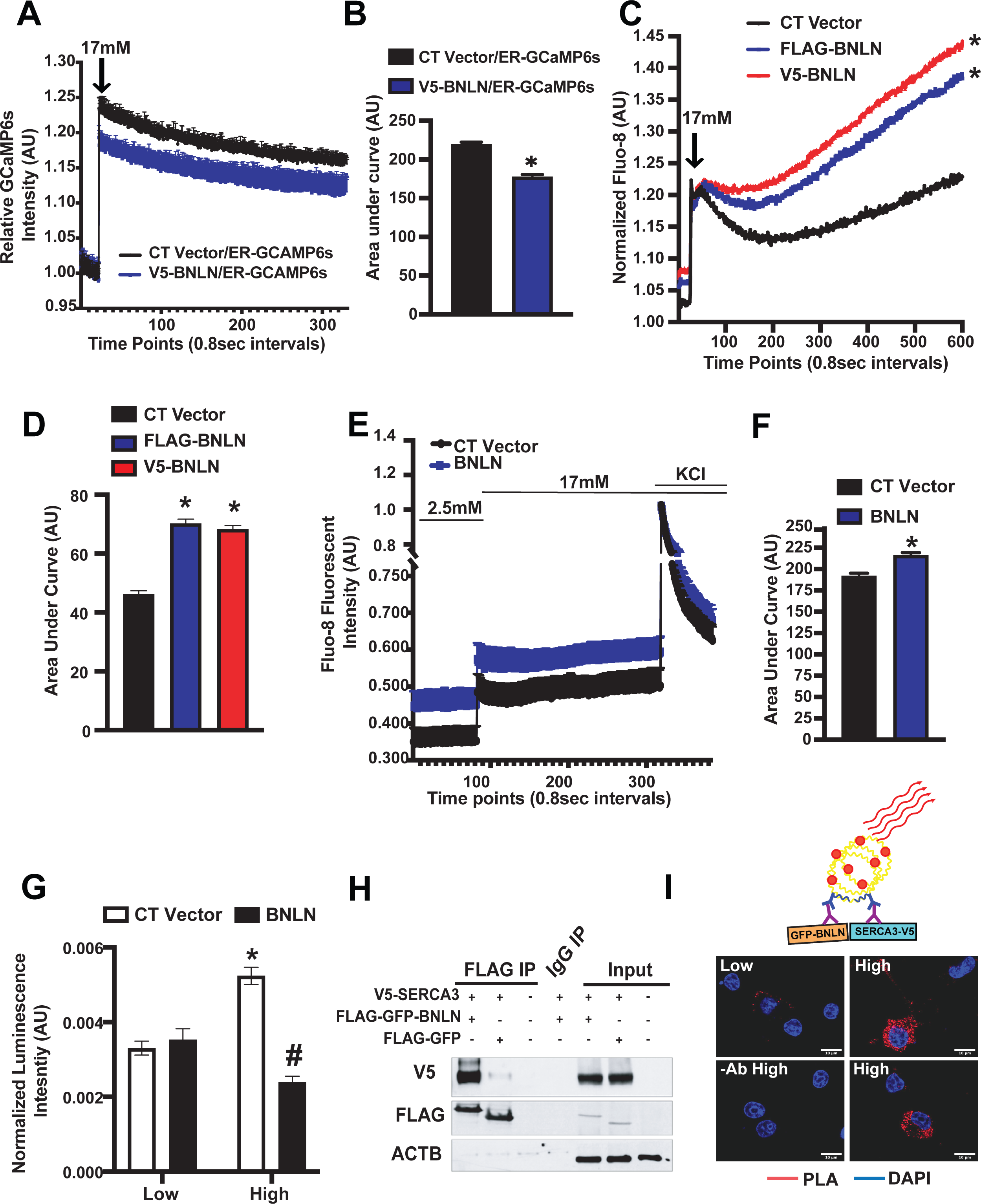
BNLN regulates glucose-induced calcium dynamics in pancreatic β-cells. **A.** Representative ER Ca^2+^ recording of INS-1 cells cotransfected with V5-BNLN and an ER calcium probe (GCaMP6s) after stimulation with high glucose (17mM) with a microplate reader at 37°C. CT; control V5 vector. **B.** Quantification of the area under a curve in (C) (n=3; experimental replicates). **C.** Representative cytoplasmic Ca^2+^ recording of INS-1 cells transfected with V5-BNLN or FLAG-BNLN after stimulation with high glucose (17mM). The fluorescent signal was measured in the emission/excitation spectrum of Fluo-8 with FLIPR Tetra System at 37°C. **D.** Quantification of the area under a curve in (E) (n=6-7; experimental replicates). **E.** Representative cytoplasmic Ca^2+^ recording of INS-1 cells overexpressed with FLAG-GFP-BNLN before (2.5 mM glucose) and after stimulation with high glucose (17mM) in the absence or presence of KCl (40mM). The fluorescent signal was measured in the emission/excitation spectrum of Fluo-8 with a microplate reader at 37°C. **F.** Quantification of the area under a curve in (E) (n=8-9, experimental replicates). **G.** ATF6LD-cluc secretion measured in INS-1 cells with BNLN overexpression in the presence of low glucose (2.5mM) or high glucose (16.7mM) for 1 −2hr at 37°C. Data were normalized to Gluc secretion (n=13-15; experimental replicates). **H.** Representative western blots of BNLN and SERCA3 in HEK293T cells transfected with the indicated constructs for 24hrs. **I.** PLA assay for SERCA3 interaction with BNLN in INS-1 cells cultured in low glucose (2.5 mM) or challenged with high glucose (17 mM). Scale bar: 10μm. -Ab: no antibody control. A model of PLA assay is shown on top of the panel. V5-SERCA3 and FLAG-GFP-BNLN were recognized by primary antibodies and secondary antibodies coupled with connector oligos. Data are shown as means ± SEM. * indicates statistical significance compared to the CT vector group in (B&D), and compared to treatment of low glucose in cells with same construct in (G); # indicates statistical significance compared to the CT group in cells treated with the same concentrations of glucose in (G). Statistical significance was determined by student’s t-test in (B&F), and ANOVA followed by Tukey’s multiple comparisons test in (D&G), p<0.05. AU: arbitrary unit.

Calcium level is tightly regulated to maintain ER chaperone activity and protein folding^35^. We found that BNLN reduced [Ca^2+^] in the ER of INS-1 cells (Fig. 4A&B). To determine whether the BNLN-mediated lowering [Ca^2+^]^ER^ effect alters ER function, we examined the functional impact of BNLN on ER homeostasis, using an ATF6 luminal domain (ATF6LD) *cypridina noctiluca (Cluc)* secretion reporter assay in INS-1 cells. In this assay, cells with abundant ER chaperones retain the ATF6LD-Cluc in the ER and secrete minimal luciferase, while cells with limited ER chaperone availability release the chimeric protein, promoting luciferase secretion into the medium^36^. We found that the high glucose challenge (17mM) led to a decreased ER chaperone availability (increased ATF6LD-cluc secretion) in INS-1 cells, which was ameliorated by BNLN overexpression (decreased ATF6LD-cluc secretion) (Fig. 4G). These data indicate that BNLN-mediated calcium dynamic in the ER did not impair ER function in INS-1 cells. In contrast, BNLN primed the β-cell ER against the high glucose-induced ER stress.

It has been demonstrated that micropetide exert its functions through interaction with protein complex^12,13,15,20^. To identify potential BNLN interacting protein partners in pancreatic β-cells, we performed a protein pull-down assay followed by mass spectrometry analysis, using lysates from INS-1 cells with BNLN-FLAG overexpression. GFP-FLAG overexpression was used as a control construct. One protein enriched in BNLN-FLAG pull down compared to GFP-FLAG pull-down was sarco-endoplasmic reticulum Ca^2+^-ATPases 3 (SERCA3; Supplemental Table 2), a major calcium transporter in the ER. We further validated this interaction by Co-IP-western blot analysis in HEK 293 cells with BNLN and SERCA3 overexpression (Fig. 4H). To determine the impact of glucose on the interaction of BNLN with SERCA3, we further performed a proximity ligation assay (PLA) in INS-1 cells in the presence of low (2.5mM) or high (17mM) glucose. As shown in figure 4I, a low level of interaction between BNLN and SERCA3 occurred in low glucose conditions (2.5mM), and this was elevated in the presence of high glucose (17mM). Taken together, these data indicate an important role of BNLN on modulating calcium dynamics in the pancreatic β-cells.

### BNLN enhances glucose-stimulated insulin secretion

Ca^2+^ oscillations in pancreatic β-cells are critical for their response to changes in glucose level^37^. To establish the functional role of BNLN in the pancreatic-β cell, we first measured glucose-stimulated insulin secretion (GSIS) in INS-1 cells with BNLN overexpression using adenovirus-mediated gene delivery. As shown in figure 5A, BNLN overexpression significantly increased GSIS in the INS-1 cells compared to cells overexpressed with control vector. To verify that this effect was mediated by the translated micropeptide and not the RNA transcript, we transfected INS-1 cells with the 2^nd^ ATG-deletion BNLN construct (which does not produce the micropeptide), and found that this construct did not affect GSIS (Fig. 5B). To further examine the BNLN-mediated insulin secretion, we transduced islets isolated from lean mice with the adenovirus-mediated BNLN construct. As shown in figure 5C, GSIS was significantly enhanced in Islets overexpressing BNLN.

**Figure 5.**
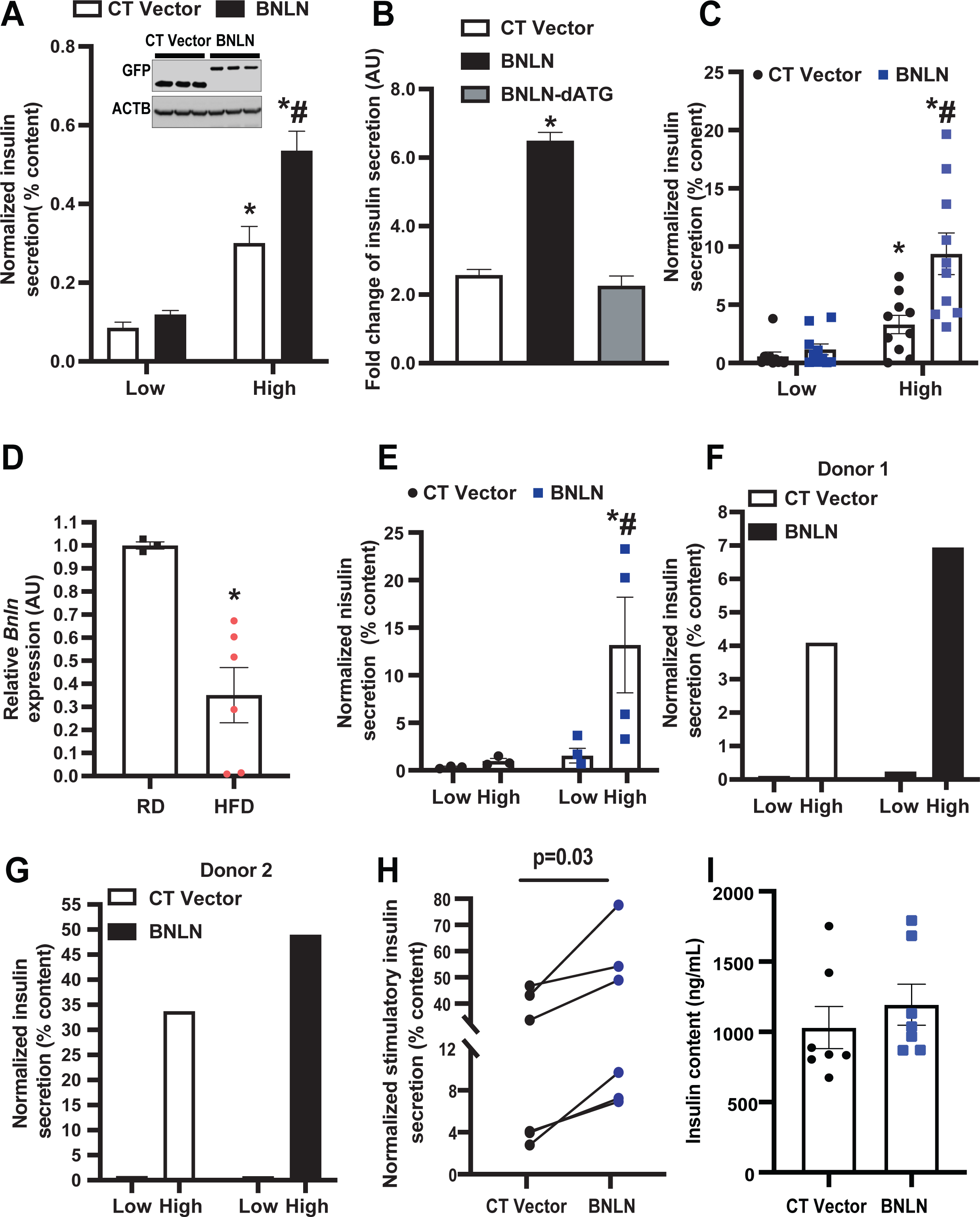
BNLN regulates glucose-stimulated insulin secretion. **A.** Glucose stimulated insulin secretion in INS-1 cells overexpressed with FLAG-GFP-BNLN in static incubation in media containing low glucose (2.5 mM) or high glucose (17 mM) for 1 hr each (n=3-4, experimental replicates). The data were normalized to insulin content. Representative western blots of BNLN expression is shown on the top of pannel. **B.** GSIS in INS-1 cells overexpressed with the indicated constructs for 48hrs in static incubation in media containing low glucose (2.5mM) or high glucose (17mM) for 1hr each (n=3-4, experimental replicates). The data were normalized to insulin content. **C.** GSIS measured in isolated islets from mice fed a RD (n = 5-10, biological replicates; 12 weeks) followed transduction of adeno-GFP (CT) or adeno-BNLN constructs (48 hrs), in static incubation in media containing low glucose (2.5mM) or high glucose (17mM) for 1hr each. The data were normalized to insulin content. **D.** Levels of mRNAs encoding for *BNLN* in islets isolated from mice fed with a RD or a HFD (12 weeks). The data were normalized to *Hprt* expression in the islets. n=3-4, biological replicates. **E.** GSIS measured in isolated islets from mice fed a HFD (n=3-4, biological replicates; 12 weeks on HFD) followed transduction of adeno-GFP (CT) or adeno-BNLN constructs (48hrs), in static incubation in media containing low glucose (2.5mM) or high glucose (17mM) for 1hr each. The data were normalized to insulin content. **F-H.** Representative GSIS measured in islets from human donors followed transduction of adeno-GFP (CT) or adeno-BNLN constructs (48hrs), in static incubation in media containing low glucose (2.5mM) or high glucose (17mM) for 1hr each. The data were normalized to insulin content. In (F), GSIS was performed on a male non-diabetic human donor of 69 years of age, 27.2 BMI, in (G), GSIS was performed on a male non-diabetic human donor of 18 years of age, 19 BMI. **I.** Insulin content measured in islets from human donors followed transduction of adeno-GFP (CT) or adeno-BNLN constructs (48hrs). Data are represented as a difference in intracellular insulin content after performing GSIS between CT and BNLN human islets. Data are shown as means ± SEM. *indicates statistical significance compared to treatment of low glucose in cells/islets with same type of construct in (A,C,E&H), to the CT vector group in (B); # indicates statistical significance compared to the CT group in cells/islets incubated with the same concentrations of glucose. Statistical significance was determined by student’s t-test in (D&H), and ANOVA followed by Tukey’s multiple comparisons test in (A&B&C&E), p<0.05. AU: arbitrary unit.

Obesity results in dysregulated GSIS in pancreatic islets. To establish the pathophysiological relevance of BNLN expression, we next assessed the expression level of *BNLN* transcripts in islets from mouse islets from lean mice and mice with diet-induced obesity (DIO). *BNLN* expression was significantly downregulated in islets from obese mice (Fig. 5D). To test whether BNLN could rescue obesity-associated islet dysfunction, we further examined GSIS in lean and obese mouse islets following *BNLN* overexpression. As shown in figure 5E, overexpression of *BNLN* significantly improved GSIS in islets from mice with DIO. Finally, to establish the functional relevance of BNLN in humans, we measured GSIS in human islets with BNLN overexpression (Supplemental Table 3). Overexpression of BNLN significantly increased GSIS in islets from non-diabetic humans with minimal effect on insulin content (Fig. 5F-I). These data strongly implicate that BNLN functions as a small peptide to modulate insulin secretion in β-cells.

## DISSCUSION

Obesity and diabetes are characterized by dysregulation of insulin secretion from pancreatic β-cells. Although lncRNAs have been implicated in this process^5–7^, thus far there is no evidence of regulatory role of lncRNA-encoded micropeptides in pancreatic β-cell function. Our study offers the first insight into the impact of micropeptides on β-cell physiology. We showed that the lncRNA *TUNAR (also known as TUNA, HI-LNC78 or LINC00617)* encodes BNLN, a 48aa micropeptide. We further demonstrate that BNLN regulates glucose-induced insulin secretion in β-cells and intact islets in part through modulating ER homeostasis.

LncRNAs regulate key biological processes through regulation of transcriptional activation, heterochromatin formation, interaction with miRNAs and other mechanisms^38,39^. However, recent studies have revealed that some transcripts identified as IncRNAs actually do contain sORFs that encode micropeptide (microprotein)^12–20^. In this study, we applied two computational programs PhyloCSF and RNAcode to prioritize the search for additional micropeptide encoding lncRNAs expressed in human pancreas. It should be noted that both PhyloCSF and RNAcode depend on the quality of whole genome sequence alignment. False negatives could be resulted from poor or incorrect whole genome sequence alignment. Among the predicted micropeptide encoding lncRNA, we choose one lncRNA, TCL1 Upstream Neural Differentiation-Associated RNA (*TUNAR*), for further characterization because it ranked at the top of both the expression level in human islets and the coding potential. *TUNAR* has been shown to be important for ESC pluripotency and neural differentiation^40,41^. Dysregulation of *TUNAR* has been implicated in the progression of Huntington’s Disease^42^ and breast cancer^5^. Notably, a recent study showed that knocking down *TUNAR* suppresses insulin secretion in a human β-cell line^5^. Multiple molecular mechanisms have been proposed underlying functions of *TUNAR* in the context of diverse biology processes. For example, it has been suggested that *TUNAR* forms an RNA–multiprotein complex to activate pluripotency genes in a β-cell line^40^. Our computational and experimental studies identified an sORF encoding BNLN, a 48aa micropeptide within the last exon of *TUNAR*. Overexpression of this micropeptide regulated GSIS, while deletion of start codon (ATG) abolished its function (Fig. 5B). However, we could not completely rule out a role for *TUNAR* lncRNA in this process. Therefore, future studies will focus on dissecting the contributions of TUNAR RNA and BNLN micropeptide on the β-cell transcriptome development and differentiation, and metabolic function.

Accumulating evidence demonstrates that transmembrane micropeptides tightly modulate intracellular and extracellular signaling cascades^43^. We found that BNLN contains a transmembrane domain at the C-terminus. To determine the sub-cellular localization, cellular function, and molecular mechanisms of BNLN in β-cells, we generated a set of epitope-tagged BNLN constructs. It is possible that epitope tags altered the signal sequences and function of BNLN, however, we did not observe functional differences between the differently tagged BNLN constructs (Fig. 4C). Further studies with specific anti-BNLN antibodies will be necessary to confirm subcellular localization of the endogenous micropeptide.

ER calcium homeostasis is required for insulin production and secretion in β-cells^26^. Ca^2+^ serves as a critical co-factor for protein chaperones and foldases needed to support high levels of protein translation and pro-insulin processing^35^. Moreover, the ER of β-cells contributes to a tight regulation of cytosolic Ca^2+^ by taking up Ca^2+^ when the cell is depolarized (cytosolic Ca^2+^ is elevated), and slowly releasing Ca^2+^ when the cell is repolarized^44^. In pancreatic islets, the ER primarily takes up cytosolic Ca^2+^ by two SERCAs: ubiquitously expressed SERCA2b, and SERCA3, which is expressed only in the islet β-cell^45,46^. While SERCA2b regulates basal [Ca^2+^]^c46^, it has been suggested that SERCA3 is responsible for replenishing ER Ca^2+^ when the cytosolic Ca^2+^ level is elevated by depolarization^47^, thus prolonging the Ca^2^ oscillation. We showed that overexpression of BNLN did not alter KCl-induced [Ca^2+^]^c^ peak, but decreased [Ca^2+^]^ER^ and increased [Ca^2+^]^c^ in response to stimulatory glucose concentrations (Fig. 4E). This is in line with a study from Ravier *et al*., which showed that SERCA3-null islets exhibited decreased ER Ca^2+^ influx, leading to an increased cytosolic Ca^2+^ oscillatory amplitude and enhanced insulin secretion^48^. Our study mainly relied on INS-1 cells as a surrogate for β-cells, and determining how BNLN modulates islet Ca^2+^ oscillation/kinetics in health and obesity/diabetes requires further studies.

We found that, in β-cells, BNLN interacts with SERCA3A (Fig. 4H&I), and decreases Ca^2+^ buffer capacity of the ER in response to high glucose (Fig. 4A&B), indicating a potential inhibitory role of BNLN on SERCA3. However, whether this is primarily the result of reduction of SERCA3A enzyme activity requires further investigation. Furthermore, sustained high-glucose elicits ER stress in the β-cell^35^. Disturbed ER homeostasis leads to impaired ER Ca^2+^ efflux, Ca^2+^-induced Ca^2+^ release (CICR), as well as dysfunction of insulin production in the β-cell. We found that the high glucose-induced ER stress was ameliorated by BNLN overexpression, while BNLN decreased ER Ca^2^” level. Further study is needed to comprehensively delineate the impact of BNLN on ER homeostasis and ER Ca^2^” leakage in β-cells under diverse ER stress conditions. Notably, glucose and amino acids promote ER Ca^2+^ uptake in the absence of external Ca^2+ 37,49^, suggesting that cytosolic Ca^2+^-binding proteins and other intracellular Ca^2+^ storage represent an important source of Ca^2+^ that can be pumped by the ER. Future study will be required to establish the functional impact of BNLN on other protein binding partners (Supplemental Table 2).

In summary, our study demonstrates that *TUNAR*, a previously annotated lncRNA, encodes a conserved 48aa micropeptide BNLN, which modulates Ca^2+^ homeostasis in β-cells, consequently regulating GSIS. This study indicates that dysregulation of BNLN in the pancreas might contribute to the obesity-associated malfunction of insulin secretion. Our study provides important insights into the molecular mechanisms that underlie the fine-tuning of glucose-induced insulin secretion by translational function of lncRNA. In addition, our study will also have a broader impact on understanding of the interactions between lncRNA biology and dysfunction of islets relevant to obesity and type 2 diabetes. It remains an intriguing possibility that insights from this study could be exploited for therapeutic interventions in obesity and type 2 diabetes.

## AUTHOR CONTRIBUTIONS

M.L. and F.S performed the experiments, analyzed the data. QW.Q, WJ.Y., ZY.Z., BY.C, D.S., YW.G., and AV.P. performed the experiments. J.S., Y.I, S.S. and LS.S. provided critical reagents and scientific suggestions on this study. HJ.C and L.Y conceived and supervised the study and wrote the manuscript. L.Y. is supported by NIH (DK108835-01A1) and American Diabetes Association (1-18-IBS-149); Z.Z. is supported by American Heart Association (19PRE34380258).

## ACKNOWLEDGEMENT

We are grateful to the Alberta Diabetes Institute Islet Core (Canada) for providing us the human islets, and the contributions of the donors whose islets were used, as well as the physicians, nurses, and researchers who procured the specimens. We also thank Dr. Michael Welsh (University of Iowa) for providing technical supports in RNA imaging; Dr. Monica Nagendran and Dr. Tushar Desai (Stanford University School of Medicine) for sharing the Proximity ligation in situ hybridization protocol with Welsh lab. We thank Dr. Timothy Ryan (Weill Cornell Medicine) for donating the ER-GCaMP6-15 to Addgene and Alyona Li for technical support with graphic illustrations. We would like thank members of the Yang, Cao, and Amendt laboratories for helpful discussion and suggestions.

## MATERIAL AND METHODS

### Prioritize putative micropeptide encoding lncRNAs in human pancreas

RNA-seq data of human pancreas islets were downloaded from EBI arrayexpress database (E-MTAB-1294^34^). RNA-seq reads were quality checked using the FastQC tool (http://www.bioinformatics.babraham.ac.uk/projects/fastqc). Low-quality and adapter sequences were trimmed using the Trimmomatic^50^. Expression of transcripts was quantified using the Salmon^51^, and estimates of transcript abundance for gene-level analysis were imported and summarized using the tximport^52^ function of the R/Bioconductor software suite^53^. Transcripts that have TPM above 1 were evaluated for their coding potential with PhyloCSF and RNAcode. We downloaded 100-species whole genome alignments (multiz100way) from the UCSC database. A custom python script was used to extract and prepare multispecies DNA sequence alignments for each transcript of protein coding (<100aa) genes and lncRNAs. We filtered for transcripts that had at least 2 species that have more than 60-nucleotide (nt) DNA sequence in the multispecies alignments. PhyloCSF and RNAcode were run with these multispecies DNA sequence alignments as input. To prioritize search of putative micropeptide encoded by lncRNAs, we ranked genes with both expression level in human islets and coding potential.

### Animals

Animal care and experimental procedures were performed with approval from the University of Iowa’s Institutional Animal Care and Use Committee. C57BL/6J mice (The Jackson Laboratory, 000664) were kept on a 12 hr light/dark cycle, and were fed the RD (7319 Teklad global diet). Mice used in generating the DIO model were placed on a 60% kcal high-fat diet (HFD; Research Diets, D12492) immediately after weaning, at 3 weeks of age.

### Human islets

Human islet study was reviewed by IRB at University of Iowa and deemed non-human study. Human islets were received from the Alberta Diabetes Institute Islet Core with reported viability and purity above 80%. Islet were cultured in CMRL1066 (Thermo Fisher Scientific, 11530037) medium with 1% human serum albumin, 1% Pen-Strep (Thermo Fisher Scientific, 15140163), and 1% L-Glutamine (Thermo Fisher Scientific, 25030164) overnight at 37°C upon arrival for recovery upon shipping. The eight donors were: age, 20-69 years; BMI, 19-33.9; Hba1c, 4.9-5.8; males; cause of death, neurological.

### Cell culture, transfection and Luciferase Assay

The rat insulinoma cells (INS-1, 832/3)^54^ were cultured in RPMI-1640 medium (Thermo Fisher Scientific, 11875119) supplemented with 10% FBS, 10mmol/L HEPES (Thermo Fisher Scientific, 15630130), 2mmol/L L-glutamine, 1 mmol/L sodium pyruvate (Sigma-Aldrich, P2256), and 50μmol/L 2-mercaptoethanol (VWR, 97064-880). Cells were transfected with DNA constructs (1ug/well in 12-well plate) using Lipofectamine 2000 reagent (Invitrogen, 11668019) in Opti-MEM medium (Thermo Fisher Scientific, 31985088). At 48 hr post transfection, the transfected cells were trypsinized (Trypsin-EDTA 0.05%; Thermo Fisher Scientific, 25300054), mixed, and re-plated in an appropriate plate in order to achieve homogeneous population across wells subsequent analyses. The HEK 293T cells (ATCC) were cultured in DMEM medium with 10% FBS, 1 % penicillin-streptomycin. Cells were transfected with DNA constructs (0.5 ug/well of 24-well plate) using Lipofectamine 3000 reagent (Thermo Fisher, L3000015) in Opti-MEM medium (Thermo Fisher Scientific, 31985-062). At 48 hr post transfection, the activities of firefly luciferase and Renilla luciferase were measured using the Luciferase Assay System (Promega, E1500) and Renilla Luciferase Assay System (Promega, E2810), separately.

### Constructs, adenovirus preparation and transduction

The BNLN constructs were generated by Sequence- and ligation-independent cloning (SLIC) method^55^. The detailed information of constructs used in each figure is list in supplementary Table2. Briefly, tRFP and Renilla luciferase-GFP expressing sequences were fused to C-terminus of human BNLN (in-frame and out-of-frame; fs1, fs2). The recombined fragments were then inserted into pCAGEN vector (Addgene, 11179). To determine transcription initial site and BNLN localization, full length human BNLN, ATGs- and CTG-deleted BNLN (dATG1, dATG2, dATG3, dCTG) were fused with Flag and cloned into pcDNA3.1(-) (Invitrogen, V79520) vector. KDEL plasmid was obtained from Addgene (114177). The HiBiT-BNLN and V5-BNLN constructs were generated by fusing HiBiT vector (Promega) and V5 sequence, respectively, in-frame with N-terminus of BNLN. The fused fragments were then inserted into pQCXIP vector (Addgene, 631516), by using NEBuilder^®^ HiFi DNA Assembly Cloning Kit (New England Biolabs, E5520S).

ER-GCaMP6-15^56^ was provided by Addgene. Adenoviruses carrying mouse *TUNA BNLN* were generated using the ViraPowerTM adenoviral expression system (Invitrogen). Adenoviruses were amplified in HEK293A cells (ATCC) and purified by CsCl gradient centrifugation. The viruses were titered using Adeno-X Rapid Titer Kit (Takaba Bio, 632250) and transduced into the cells as previously described ^57^. Cultured intact mouse or human islets were resuspended in 0.1mmol/L EGTA (Sigma-Adlrich, 234626) in serum free CRMLfollowed by transduction of indicated adenoviral constructs at 10,000 plaque-forming unit per islet equivalent (pfu/IEQ) for 1hr at room temperature before transferring to CMRL medium with 10% heat-inactivated FBS for culture overnight[]. The culture medium was replaced next day and the islets were cultured for additional 24hr before subjecting them to glucose stimulated insulin secretion (GSIS) assay.

### Nano-Glo HiBiT extracellular detection system

Ins-1 cells were transfected with FLAG-GFP or HiBiT-BNLN as described above and seeded in an opaque 96-well plate(Fisher Scientific, 08-771-26) at ~15,000 cells/well in the culture medium. The experiment was carried out according to the manufacturer’s protocol (Promega, N2422). Briefly, the transfected cells were incubated with Nano-Glo HiBiT extracellular buffer, LgBiT protein, and Nano-Glo HiBiT extracellular substrate, and luminescence was detected after 5-10min using a luminescence microplate reader (SpectraMax, Molecular Devices, USA). The expression of extracellular HiBiT-BNLN was normalized to the total intracellular pool of HiBiT-BNLN by lysing the cells and measuring luminescence with Nano-Glo HiBiT lytic detection system (Promega, N2422), following the protocol.

### Western Blotting and immunoprecipitation

Cells were lysed with a lysis buffer containing 0.4% of NP-40, protease inhibitors cocktail (Sigma-Aldrich, P8340), and Na3VPO4 (Sigma-Aldrich). Protein concentration was determined by Pierce BCA kit (Thermo Fisher Scientific, 23225) and samples were subjected to SDS-polyacrylamide gel electrophoresis, as previously described (48). For immunoprecipitation assay, cells were lysed in radioimmunoprecipitation assay (RIPA) buffer (50mM Tris, 150mM NaCl, 1% NP40, 0.25% sodium deoxycholate). 1mg proteins were immunoprecipitated with protein A/G magnetic Beads (ThermoFisher, 88802) conjugated with anti-IgG (Cell Signaling Technology, 2729), anti-Flag (Thermo Fisher, A36797) and eluted by 4× lituium dodecyl sulfate (LDS; Invitrogen, NP0008). Membranes were immunoblotted with anti-V5 (ab15528), anti-FLAG and anti-beta Actin (ab8227) at a 1:1000 dilution. Secondary antibodies were horseradish peroxidase-conjugated goat-anti-mouse-IgG (Santa Cruz Biotechnology, sc-2005), horseradish peroxidase-conjugated mouse-anti-rabbit-IgG (Santa Cruz Biotechnology, sc-2357) at a 1:10000 dilution. Signal was detected using the ChemiDoc Touch Imaging System (BioRad), and densitometry analyses of western blot images were performed by using the Image Lab software (BioRad).

### Proximity ligation *in situ* hybridization (PLISH)

*In situ* hybridization was performed as previously described^58^ with some modifications. Briefly, human islets were fixed and sectioned following the typical formalin-fixed paraffin embedded tissue processing protocol. The sections were then incubated in citrate-based target unmasking solution (Vector laboratories, H-3300) with 0.05% lithium dodecyl sulfate (Sigma, L4632) at 65°C for 30 min. The slides were then incubated with 0.05 mg/ml pepsin (Sigma-Aldrich, P6887) in 0.1M HCl for 10 min at 37 °C, and processed with the typical proximity ligation *in situ* hybridization^58^. RNA sequences that were targeted by short-paired hybridization probes were: Human INS-1: CTGGTGGAAGCTCTCTACCTAGTGTGCGGGGAACGAGGCT;Human INS-2: GCTGGAGAACTACTGCAACTAGACGCAGCCCGCAGGCAGC; HumanGCG-1: TGGACT CCAGGCGTGCCCAAGATTTTGTGCAGTGGTTGAT; Human GCG-2: AGATGAACACCATTCTTGATAATCTTGCCGCCAGGGACTT; Human GCG-3: TCTTCACAACATCACCTGCTAGCCACGTGGGATGTTTGAA; Human BNLN: TCTTGGCAATGCTGGGGATTATCGGGACCATTCTGAACCT; Human BNLN: AGCA GGCTTGACCCGCACATACCACCCAATCAAATGCACC’; Human BNLN: ATTCCAGATCGCTGACAGAT ATCACATATTTGAAAAGATG; Human BNLN: CTTCACTTGACGAGCTATTTAGTGGAAAAACCACAGG CGC; Human BNLN: GCAGACCTTAGATGCACCCTATCTTTACTGAGAATTATGC; Human BNLN: TTTGATTTTAGCGGTCATGTACCGCGAGAGTTGGGAAGAA.

### Immunocytochemistry and Confocal microscopy

INS-1 cells were then fixed in 4% paraformaldehyde for 10min at room temperature, blocked in 5% goat serum, incubated with Hoechst33342 (Thermo Fisher Scientific, H3570), and mounted with a glass coverslip. For subcellular organelle localization, Lystroacker™ Red DND-99 (Thermo Fisher Scientific, L7528), Mitotracker™ Red CMXRos (Thermo Fisher Scientific, M7512), ACAA1 antibody (Gene Tex, GTX114229), RFP-KDEL plasmid, and TGN38 antibody (Novus, NB300-575) were used to detect lysosomes, mitochondria, peroxisomes, the ER, and Golgi respectively. Images were acquired using a Zeiss 700-point scanning confocal microscope with a 63x lens with oil (Zeiss, Oberkochen, Germany). Images were analyzed by Fiji software. During analysis, single Z-stacks were processed and images from different channels were merged. For PLISH assays, stack images were taken using the 60x lens of Olympus microscopy (FV3000, confocal laser scanning microscope).

### Quantitative real-time RT-PCR

Total RNA was isolated using the Trizol reagent (Invitrogen, 15596026) and reverse transcribed into cDNA using a PrimeScript RT RT Master Mix (Takara RR036A). Quantitative real-time RT-PCR analysis was performed using TB Green Premix Ex Taq (Clontech, RR420A) and following primers: *BNLN*-F: 5’-TTC TCC TCG CCT TCC TGC-3’;*BNLN*-R: 5’-CAT CTT GGT TGC AAA TGT CT-3’; *ACTB*-F: 5’-CTC TTC CAG CCT TCC TTC-3’; *ACTB*-R: 5’-ATC TCC TTC TGC ATC CTG TC-3’; *Hprt-F:* 5’-CAG TCC CAG CGT CGT GAT TA-3’; *Hprt-R:* 5’-GGC CTC CCA TCT CCT TCA TG-3’; *Bnln-F:* 5’-GGA GAA TGA GGC AGG AAC CC-3’; *Bnln*-R: 5’ACA CGC TCT CTT CCT TGC TC-3’.

### Calcium imaging

#### [Ca^2+^]^c^ measurement

INS-1 832/3 cells were transfected with FLAG-GFP or FLAG-GFP-BNLN. 24hr after transfection, the Ins-1 cells were plated at ~25,000 cells/well in a HTB9-coated Greiner Sensoplate™ glass bottom 96-well plate (Millipore Sigma). Prior to imaging, the cells were loaded with Screen Quest™ Fluo-8 no wash calcium red cye (AAT Bioquest, 36314) indicator in the DMEM (Thermo Fisher Scientific, A1443001) culture medium supplemented with 2.5mM glucose and 0.1% BSA for 1hr at 37°C. A fluorescent microplate reader (CLARIOstar PLUS, BMG Labtech, Germany) was used to inject glucose (final concentration of 20mM) to cells, and to acquire fluorescent signal every 0.6 sec for 10min.

For measurement of [Ca^2+^]^c^ using Fluorescent Imaging Plate Reader(R) (FLIPR) Tetra^®^ System (Molecular Devices, USA), INS-1 cells were transfected with pQCXIP, FLAG-BNLN, or V5-BNLN (1ug DNA/well) in a 12-well plate (3 wells/each DNA construct) with Lipofectamine 2000, as described above. A suspension of Ins-1 cells was pipetted in a 384-well culture black microplate with optic bottom plate (Greiner-Bio, 781090) at ~17,000 cells/well in 25uL of the low glucose DMEM culture medium. 25uL of Fluo-8 calcium dye was used added to the cell suspension. The plate was incubated at 37° for 30min, followed by another 30min incubation at room temperature in dark. Meanwhile, a 384-well compound plate (Costar, 3657) was loaded with low glucose (27.5mM), high glucose (181.5mM), and potassium chloride as a positive control (220mM). The FLIPR was utilized to transfer the solutions from the compound plate into the black plate and to read the Ca^2+^ signal. The basal signal was measured for 20 seconds every second, then the solutions were added (5uL/each), and the response signal was read for 10min every 0.8sec.

#### [Ca^2+^]^ER^ measurement

INS-1 cells were co-transfected with ER-GCaMP6s and pQCXIP or V5-BNLN using Lipofectamine 2000, as described above. Next day, the cells were plated at ~25,000 cells/well in a HTB-9 coated plate, as described above. 24hours later, the cells were subjected to treatment with DMEM containing 2.5mM glucose at 37° for 1hr. Basal signal was acquired in the GFP excitation/emission spectrum using the fluorescent microplate reader as described above every 0.8sec for 1min. Then a high glucose solution (17mM final concentration) was injected and the fluorescent signal was collected for 5min.

### Murine islet isolation

The islets were isolated as previously described^59^. Briefly, lean C57BL/6J male mice (12-20 weeks of age) and mice with diet-induced obesity (12-16 weeks of HFD) were anesthetized with isoflurane, and the pancreas was perfused with HBSS containing type V collagenase (0.8mg/mL) via the common bile duct. The pancreas was then removed and digested at 37°C for 10-12 min with physical agitation to release islets. Islets were further washed with RPMI-1640 with 1 % FBS and purified on Histopaque 1077 and 1119 gradients Sigma-Aldrich, 10771, 11191). The islets were then collected, washed, and cultured in RPMI-1640 supplemented with 10% FBS, 1% pen-strep, and 1% HEPES in 30mm dishes at 37°C.

### Glucose stimulated insulin secretion (GSIS)

#### For GSIS in murine islets

30-40 islets of similar size were picked and placed in separate wells in a plastic 12-well plate (Fisher Scientific, 07-200-82). The islets were rinsed with 1mL of Secretion Assay Buffer (SAB) adjusted to pH7.2, containing 0.114mol/L sodium chloride (Sigma Aldrich, S9888), 0.47mmol/L potassium chloride (Sigma Aldrich, P9333), 0.12mmol/L potassium dihydrogen phosphate (Sigma Aldrich, 1.04871), 0.116mmol/L magnesium sulfate (Sigma Aldrich, M2643), 20mmol/L HEPES (Thermo Fisher Scientific, 15630130), 2.5mmol/L calcium chloride (Sigma Aldrich, 449709), 0.2% bovine serum albumin, and 0.2% sodium bicarbonate (Sigma Aldrich, S5761)^59^. The islets were then equilibrated in SAB containing low glucose (2.5mmol/L) for 1hr at 37°C. Next, the islets were incubated in fresh SAB with low glucose for additional 1-2 hours at 37°C. The supernatant was collected for measuring basal insulin secretion. The islets were then incubated with SAB containing high glucose (16.7mmol/L) for 1-2 hours at 37°C. The supernatant was collected for measuring glucose-stimulated insulin secretion. The islets were lysed in acidified ethanol and insulin content was measured. Secreted insulin and insulin content were analyzed with STELLUX^®^ Chemi Rodent Insulin ELISA (ALPCO, 80-INSMR-CH01), according to the manufacturer’s protocol.

#### For GSIS in Human islets

30-40 islets of similar size were picked and subjected to GSIS similar to murine islets. Secreted insulin and insulin content were analyzed with STELLUX^®^ Chemi Human Insulin ELISA (ALPCO, 80-INSHU-CH01), according to the manufacturer’s protocol.

#### GSIS in Ins-1 cells

100% confluent cells were rinsed with SAB and equilibrated with low-glucose-SAB for 1-2 hours at 37°C. Fresh SAB containing low glucose was added to the cells for additional 1-2 hours at 37°C. The supernatant was then collected for measuring basal insulin secretion. SAB containing high glucose was added to the cells and incubated for additional 1−2 hours at 37°C before supernatant collection. The cells were lysed and protein was extracted for insulin content. Secreted insulin and insulin content were analyzed with STELLUX^®^ Chemi Rodent Insulin ELISA (ALPCO), according to the manufacturer’s protocol.

### Statistics

The data are presented as mean ± standard error of the mean or standard deviation as noted; *n* represents the number of individual experiments; *N* represents the number of human donors or mice. Statistical differences of numeric parameters between two groups were determined with Student’s *t*-test. For statistical analysis of differences between multiple groups, one way and two-way ANOVA were applied, followed by post-hoc Tukey’s test. The statistical analyses were performed in Prism.

## Supplementary Material

**Supplemental Figure 1. Expression of TUNAR in various human tissues.** Violin plot of *TUNAR* RNA expression across various human tissues. Expression values are measured by log10(TPM+1), data from GTEX database.

**Supplemental Figure 2. Characterization of BNLN. A.** Ribosomal (Ribo-seq) profiling track from GWIPS-viz genome browser indicating translation of sORF at the begining of the last exon. **B.** TMHMM algorithms predict a transmembrane domain at the C-terminus of BNLN.

**Supplemental Figure 3. Conservation of BNLN across vertebrates species.** 88 species out of 100 vertebrate species (UCSC multiz100way) contain homologous sORFs. Multi-species alignment of *in silico* translated putative BNLN protein sequence.

