## Supplementary figures and images for "A putative long noncoding RNA-encoded micropeptide maintains cellular homeostasis in pancreatic β-cells"

### Supplemental Figures

#### Supplemental Figure 1

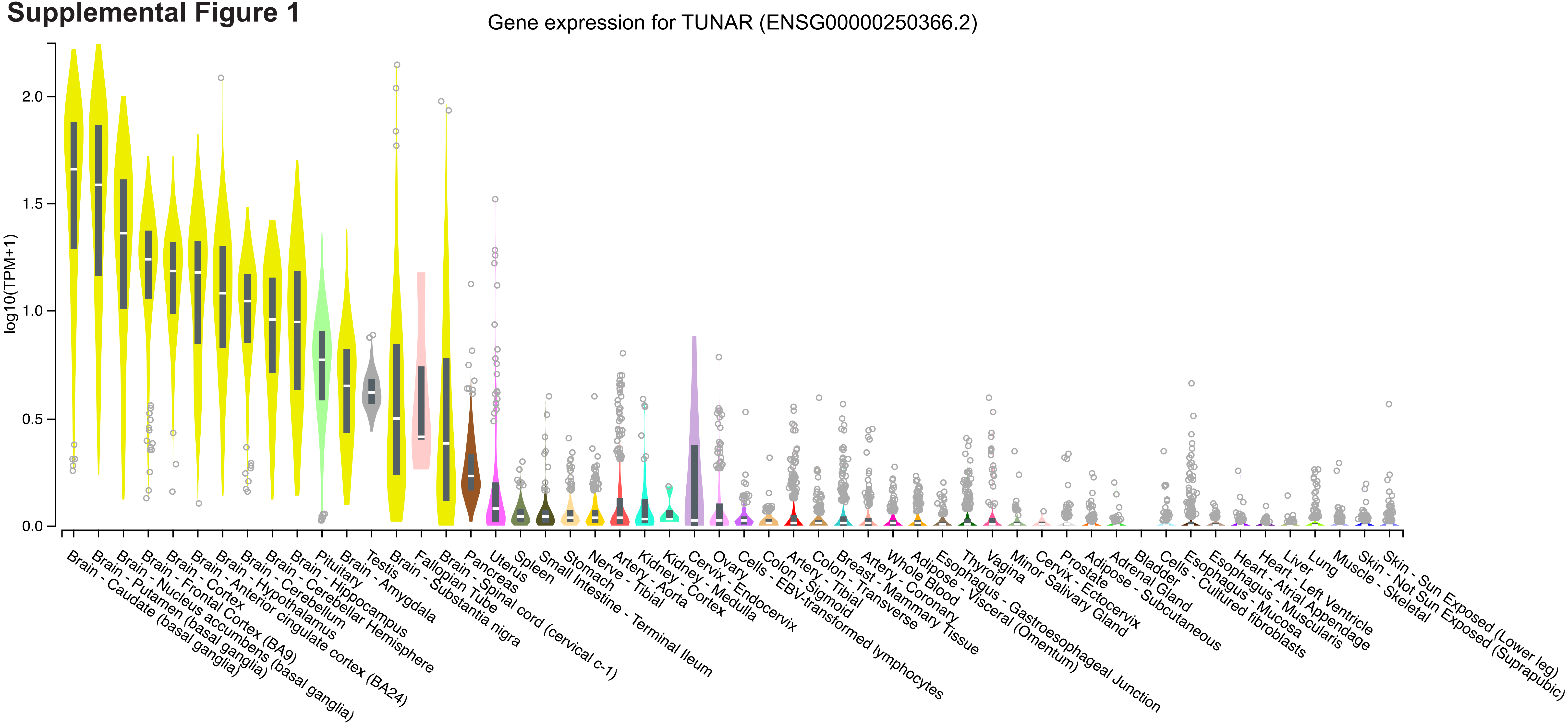

#### Supplemental Figure 2

**A**

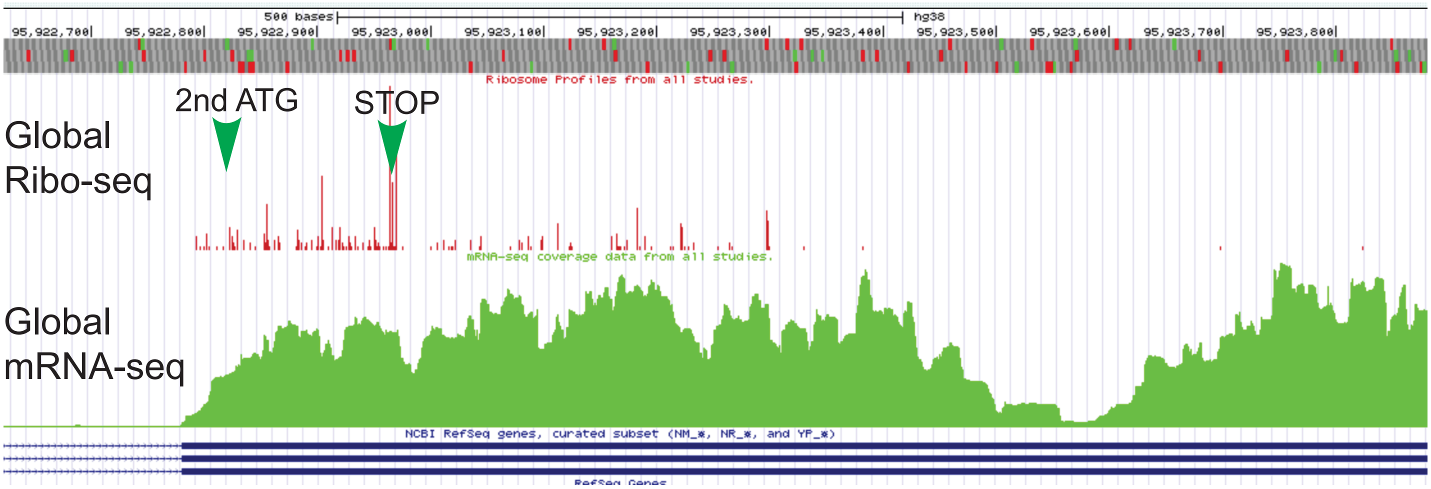


**B**

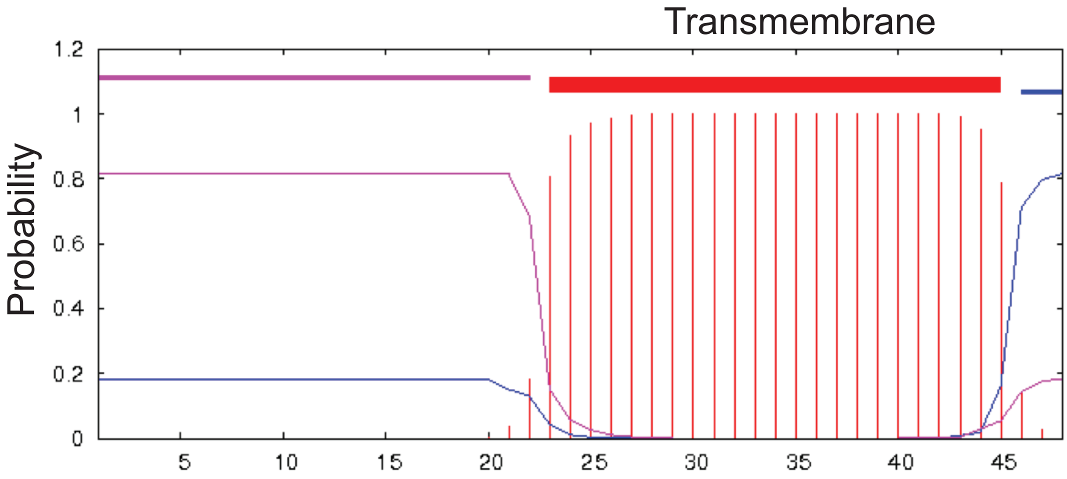

#### Supplemental Figure 3

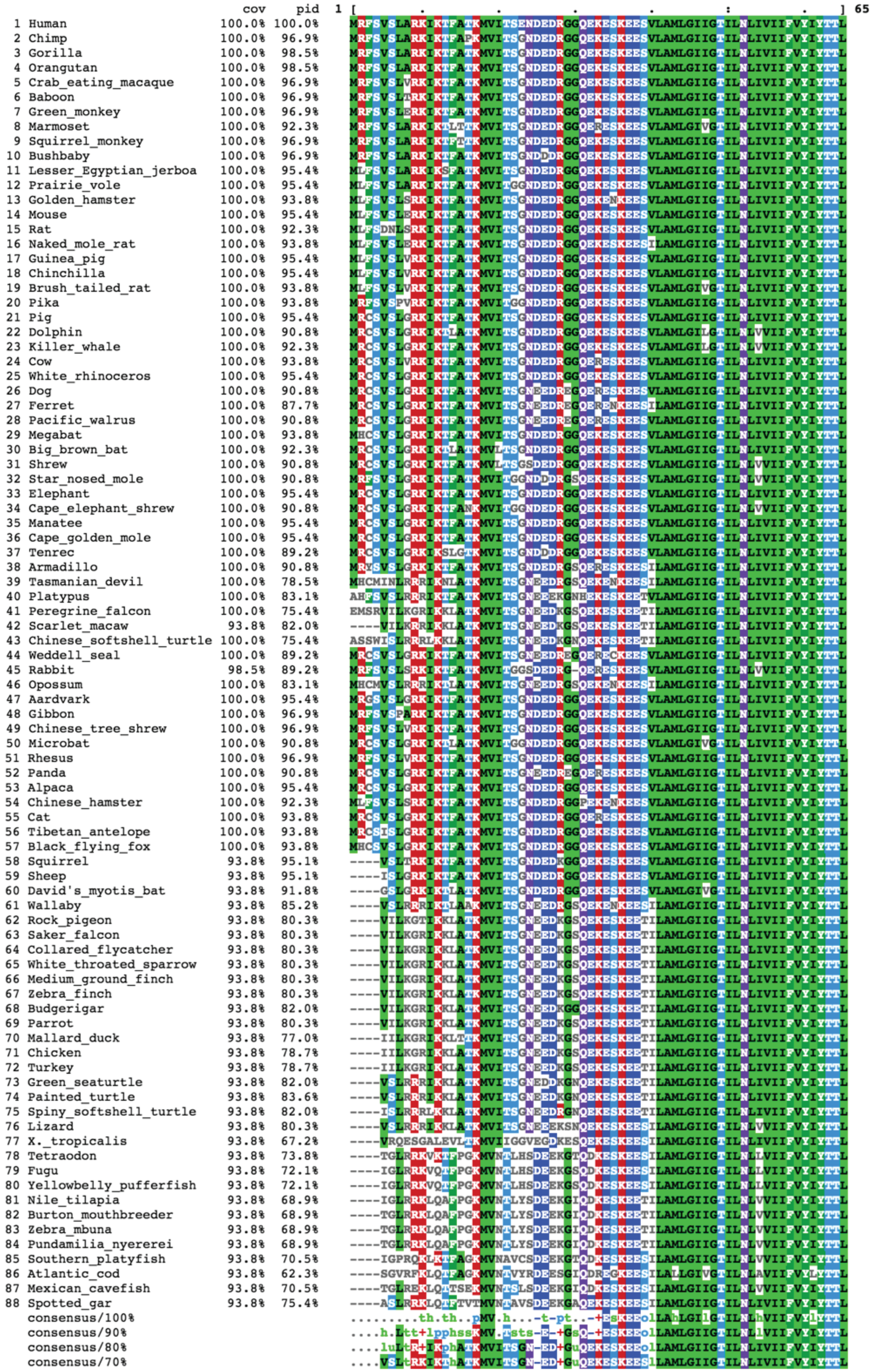
